# HSP70 binds to specific non-coding RNA and regulates human RNA Polymerase III

**DOI:** 10.1101/2023.04.10.536203

**Authors:** Sergio Leone, Avinash Srivastava, Barbara Hummel, Lena Tittel, Fernando Aprile-Garcia, Andrés Herrero-Ruiz, Prashant Rawat, Anne E. Willis, Ritwick Sawarkar

**Affiliations:** MRC, University of Cambridge, Cambridge, UK; Max Planck Institute of Immunobiology and Epigenetics, Freiburg, Germany

**Keywords:** Molecular chaperones, HSP70, RNA-binding proteome, RNA polymerase III, tRNA

## Abstract

Molecular chaperones are critical for protein homeostasis and are implicated in several human pathologies such as neurodegeneration and cancer. While the binding of chaperones to nascent and misfolded proteins has been studied in great detail, the direct interaction between chaperones and RNA has not been systematically investigated. Here we provide the evidence for widespread interaction between chaperones and RNA in human cells. We show that the major chaperone Heat-Shock Protein 70 (HSP70) binds to non-coding RNA transcribed by RNA Polymerase III (Pol III) such as tRNA and 5S rRNA. Global chromatin profiling revealed that HSP70 binds genomic sites of transcription by Pol III. Detailed biochemical analyses showed that HSP70 facilitates transcription of its target non-coding RNA by binding to Pol III. Thus our study uncovers an unexpected role of HSP70-RNA interaction in the biogenesis of a specific class of non-coding RNA with wider implications in neurodegeneration and cancer.

## INTRODUCTION

Molecular chaperones such as Heat Shock Proteins (HSP) 70, 90, and small HSPs play a critical role in the folding of nascent polypeptides, stability of folded proteins and refolding or degradation of misfolded proteins. The direct interaction of chaperones with proteins, from their synthesis to degradation, controls protein homeostasis and the functionality of the cellular proteome (Rosenzweig et al., 2019; Vabulas et al., 2010). Proteotoxic stress typically leads to induction of chaperone proteins such as HSP70 and HSP90 to mitigate the proteome damage caused by stress. Age-dependent decrease in expression and malfunction of chaperones appears to be a major driver of neurodegenerative diseases (Brehme et al., 2014; Gidalevitz et al., 2006). The dependence of cancer cells on chaperone activity has led to successful pharmacological targeting of chaperones in cancer treatment (Antonova et al., 2019; Shevtsov et al., 2019; Wen et al., 2014). Thus, chaperones play a pivotal role in cellular physiology, directly influencing organismal health and disease.

While the binding of chaperones to polypeptides has been extensively investigated, the direct interaction between chaperones and polynucleotides such as DNA and RNA has not been studied in detail. Chaperones are known to be implicated in several biological processes involving RNA, ranging from their biogenesis (Hummel et al., 2017; Sawarkar et al., 2012) and translation (Willmund et al., 2013) to ribonucleoprotein complexes formation (Iwasaki et al., 2010). Interestingly, recent data suggest that RNA may itself possess chaperone activity and may further facilitate protein refolding by the prokaryotic HSP70 chaperone system (Docter et al., 2016). A first hint of chaperone interaction with RNA was reported for the procaryotic chaperone GroEL (Georgellis et al., 1995). In the context of cell extracts, HSP70 was shown to bind AU-rich sequences present at the 3’-untranslated region (3’-UTR) of several mRNA and to regulate their stability (Henics et al., 1999; Kishor et al., 2013, 2017; Wilson et al., 2001; Zimmer et al., 2001). The interaction between HSP70 and mRNA is thought to influence mRNA stability by mechanisms that are not fully understood. It has been suggested that HSP70 recruitment to rDNA genomic loci occurs by direct binding to the newly synthetized intergenic rRNA in response to stresses such as heat shock or acidosis (Audas et al., 2012; Jacob et al., 2013). Furthermore, the yeast homologue of HSP70 (Ssa2p) has been described to interact with tRNA in the cytoplasm and mediate tRNA import into the nucleus under starvation conditions (Takano et al., 2015).

HSP70 is an ATP-dependent chaperone consisting of an N-terminal nucleotide binding domain (NBD) linked to a C-terminal substrate binding domain (SBD). HSP70 chaperone activity is based on an allosteric mechanism that couples ATP hydrolysis in the NBD to substrate binding by the SBD in a cycle of association and release that prevent substrate aggregation and promote folding (Rosenzweig et al., 2019). Both NBD and SBD can be targeted by small molecule inhibitors that reduce HSP70 activity and have been tested for cancer treatment in several clinical trials (Shevtsov et al., 2019). There is no known RNA-binding domain in HSP70 or any other chaperone protein, raising the question if chaperones directly bind to RNA in intact cells or if chaperone-RNA interaction is an artifact of promiscuous interactions in cell extracts. Recent unbiased studies have identified several proteins that directly interact with RNA but do not have a canonical RNA-binding domain (Baltz et al., 2012; Bao et al., 2018; Castello et al., 2012; Gebauer et al., 2020; He et al., 2016; Huang et al., 2018; Queiroz et al., 2019). These studies make use of UV-induced crosslinks between proteins and their interacting RNA in intact cells, suggesting the possibility of non-canonical ways of RNA-protein interactions in cells. We referred to the datasets provided by these studies to identify and experimentally validate chaperone-RNA interactions in human cells. By using the HSP70 family member HSPA1A as an exemplary chaperone, we provide a mechanism by which chaperone-RNA interaction contributes to cellular physiology.

## RESULTS

### The RNA-binding chaperome

Recent proteome-wide studies have identified a large number of eukaryotic proteins that directly interact with RNA in cells (Baltz et al., 2012; Castello et al., 2016; Gebauer et al., 2020; Queiroz et al., 2019). RBPbase, a comprehensive database that integrates several independent datasets, reports 4257 RNA-binding proteins in the human proteome (https://rbpbase.shiny.embl.de). To systematically investigate if chaperones have been found in the published proteome-wide studies of RNA-binding proteins, we focussed on 332 proteins that constitute the human chaperome (Brehme et al., 2014). 151 chaperome proteins were reported as RNA-binding proteins (Figure 1A). 99 out of these 151 proteins were found in the interactome of poly(A) RNA (Figure 1A), implicating around one third of the chaperome in direct interaction with mRNA. At least one member of each chaperome family was identified as an RNA-binding protein, with 33 chaperome proteins detected in more than 10 independent datasets (Figure 1B). There was a large overlap among several distinct studies generated using four human cell lines, with 31 chaperome proteins identified in all 23 datasets (Figure 1C). We focussed on HEK293 as data from this cell line reported the largest number of RNA-binding chaperome proteins (Figure 1C, S1A). There was no correlation between expression levels of chaperome proteins and their detection as RNA-binding proteins in HEK293T (Figure S1B). This observation rules out the possibility that highly expressed chaperones were likely contaminants in mass spectrometry-based detection of RNA-binding proteins. Thus, unbiased proteome-wide studies suggest that about half of the chaperome proteins directly bind to RNA in human cells.

**Figure 1.**
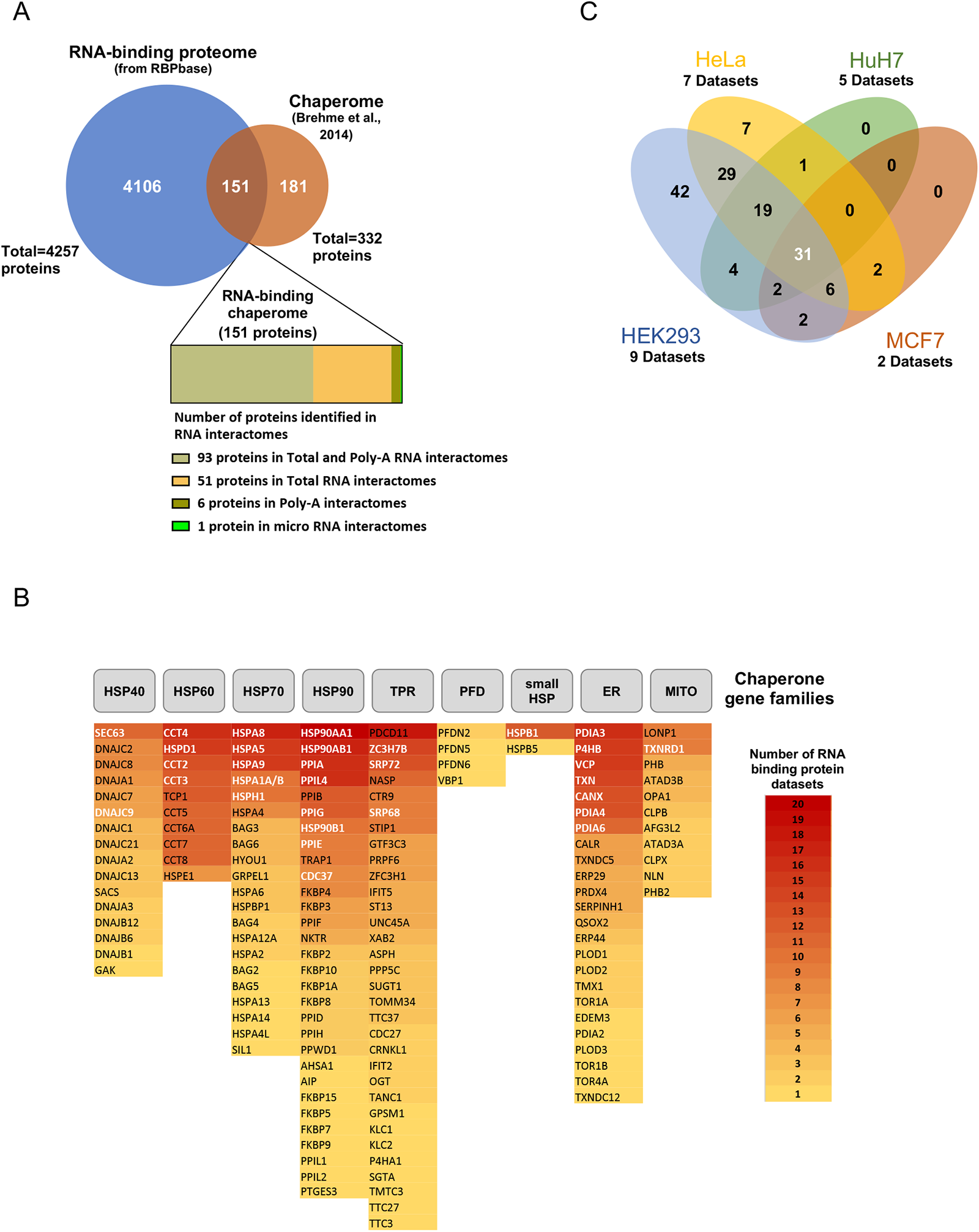
The RNA-binding Chaperome. **(A)** A Venn diagram showing the overlap between the human chaperome (from Brehme et al. 2014) and the RNA-binding proteome (from RBPbase; https://rbpbase.shiny.embl.de/, a collection of 30 independent human RNA-binding proteins datasets). The inset below the Venn diagram shows the number of RNA-binding chaperome proteins identified in Poly-A, microRNA and total RNA interactome studies. **(B)** List of the 151 RNA-binding chaperome proteins shown in (A) sorted into chaperone families (TPR: tetratricopeptide repeat (TPR)-domain-containing protein; PFD: prefoldin; ER: endoplasmic reticulum specific chaperome proteins; MITO: mitochondria specific chaperome proteins). The boxes containing the protein name are color-coded and sorted in descending order based on the number of independent datasets in which they were identified as RNA-binding proteins. The 31 chaperome proteins common to the four cell lines shown in (C) are written in white font. Number of datasets reporting HSPA1A and HSPA1B are merged since the two proteins have 100% amino acid identity and are not distinguishable by mass spectrometry analyses. **(C)** A Venn diagram showing the overlap between RNA-binding chaperone proteins independently identified in distinct human cell lines. The number of datasets analysed per cell line is indicated.

To experimentally validate the proteome-wide studies, we expressed selected chaperome proteins with C-terminal FLAG-hexa-histidine-biotin (3xFlag-HBH) tag (Aktaş et al., 2017; Maticzka et al., 2018) in HEK293T cells. UV-crosslinking was used in intact cells to covalently link chaperones with their interacting RNA. Tandem affinity purification of the chaperome proteins using hexa-histidine and biotin tags with stringent washes allowed us to enrich protein-RNA complexes. We visualized chaperome proteins in an autoradiograph after radiolabelling the cross-linked RNA using ^32^P-ATP and polynucleotide kinase (PNK; Figure 2A). RNase I-sensitive radioactive signal above and at the size of the tagged chaperome protein represented covalent protein-RNA complex, as seen for all tested chaperome proteins (Figure 2B-E). RNA-binding proteins such as HNRNPC and GAPDH (Singh and Green, 1993) served as positive controls for PNK assay (Figure 2F). Thus, our experiments provide direct evidence that several chaperome proteins directly bind to RNA, raising the question about the role of chaperones in RNA synthesis, stability and/or function.

**Figure 2.**
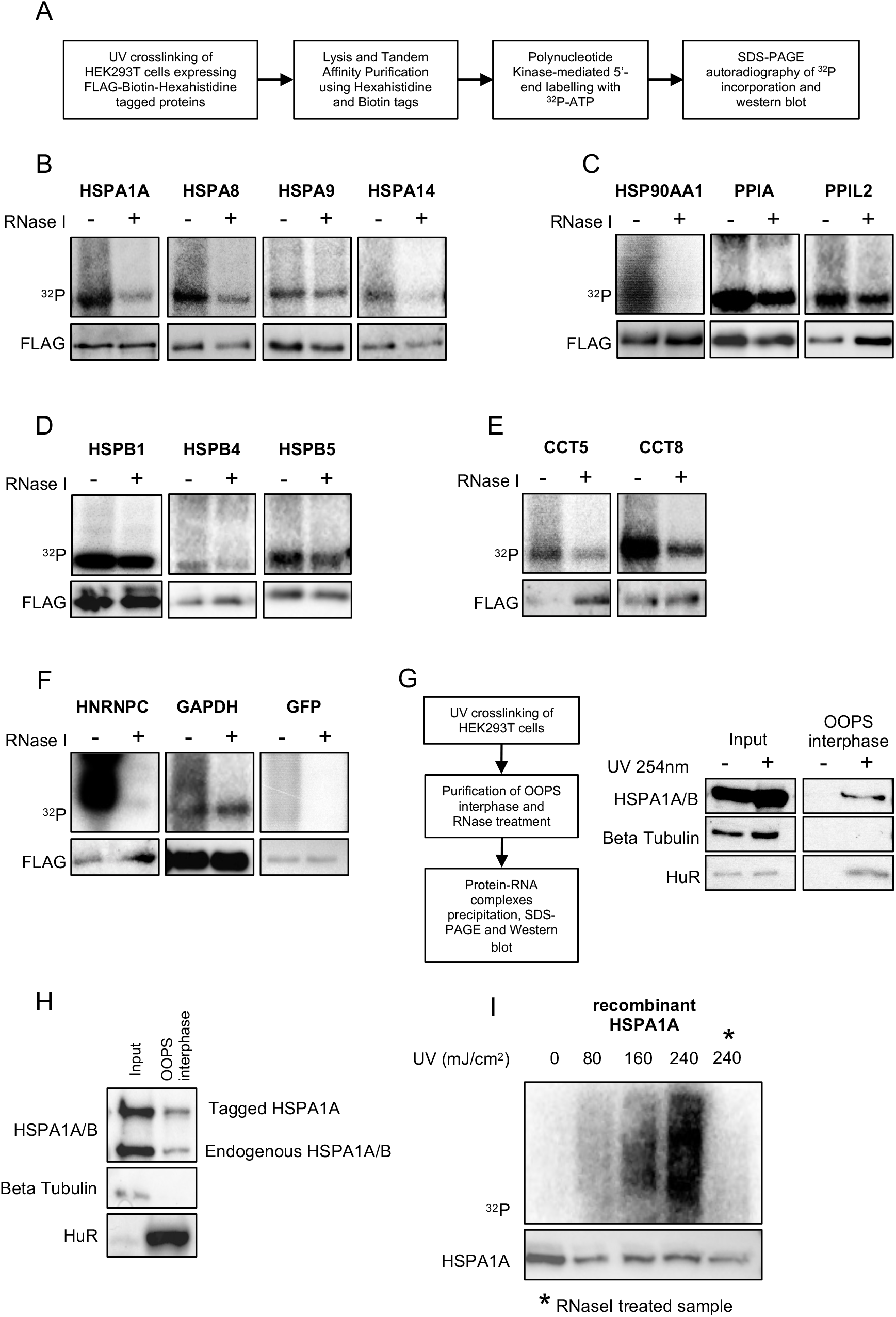
HSPA1A binds to RNA in cells and *in vitro*. **(A)** Scheme showing the experimental workflow of polynucleotide kinase (PNK) assay used to investigate direct protein-RNA interactions. **(B-F)** Validation of RNA-binding ability of chaperone proteins belonging to the HSP70 (B), HSP90 (C), small HSP (D) and HSP60 (E) families, along with control proteins (F) using the PNK assay outlined in (A). Upper panels show SDS-PAGE autoradiograph of ^32^P incorporation. Treatment with RNase I after PNK reaction is indicated. Lower panels show western blot of the affinity purified 3xFLAG-Biotin-Hexahistidine tagged chaperone proteins detected with anti-FLAG antibody. **(G)** Scheme of Orthogonal Organic Phase Separation (OOPS) experimental workflow (left). Western blot of OOPS interphase purified proteins detected with indicated antibodies (right). **(H)** Western blot of OOPS experiment performed using HEK293T cells expressing 3xFLAG-Biotin-Hexahistidine tagged HSPA1A. **(I)** PNK-assay performed using recombinant HSPA1A incubated with purified total RNA from HEK293T cells. SDS-PAGE autoradiograph of ^32^P-labelled RNA-protein complexes (upper panel) and western blot with anti-HSPA1A/B antibody detection (lower panel). The amount of UV irradiation used in the PNK assay is indicated in mJ/cm^2^. * denotes the sample treated with RNase I after PNK reaction.

### HSPA1A binds to RNA in cells and *in vitro*

To identify the significance of chaperone-RNA interactions, we focussed on HSP70 family member HSPA1A which showed strong interaction with RNA in PNK assays (Figure 2B). HSPA1A is induced during heat-shock, however the protein is present at high levels without heat-shock in HEK293T cells allowing us to dissect the role of HSPA1A-RNA interaction in unstressed cells. To verify the direct binding of endogenous HSPA1A to RNA we applied the orthogonal organic phase separation (OOPS; Queiroz et al., 2019) method. This protocol allows the enrichment in the acidic guanidinium-thiocyanate-phenol-chloroform (AGPC) interphase of protein-RNA adducts generated by UV-mediated covalent crosslinking (Figure 2G). We performed OOPS using HEK293T cells exposed to 254 nm UV light. UV-crosslinking resulted in the partitioning of endogenous HuR, a well-known RNA binding protein, as well as HSPA1A/ B, in the AGPC interphase. The negative control beta-tubulin was not found in the interphase (Figure 2G). Commercially available antibodies do not distinguish between the two identical proteins HSPA1A and HSPA1B. Hence, we resorted to an ectopic expression of 3xFlag-HBH tagged HSPA1A. We found endogenous HSPA1A/B as well as tagged HSPA1A to partition in the AGPC interphase (Figure 2H), implying the formation of UV-crosslinked HSPA1A-RNA adducts and confirming the binding ability of HSPA1A to RNA in intact cells.

To investigate if HSPA1A can directly bind to RNA outside the cellular context, we performed *in vitro* PNK assay by UV-crosslinking recombinant HSPA1A with total RNA purified from HEK293T cells. The *in vitro* approach confirmed the RNA-binding ability of HSPA1A as shown by the accumulation of radioactive signal that was proportional to the amount of UV radiation and sensitive to RNase I (Figure 2I). In summary, using three independent assays and conditions, our data confirm the ability of HSPA1A to directly bind to RNA in human cells.

### HSPA1A binds to non-coding RNA

To identify the RNA species bound by HSPA1A in cells, we took advantage of the recently developed method called FLASH-seq (fast ligation of RNA after some sort of affinity purification for high-throughput sequencing; Figure 3A) (Aktaş et al., 2017). UV-crosslinked protein-RNA complexes were enriched by tandem affinity purification of HSPA1A or GFP as a negative control. Deep sequencing of libraries made from enriched RNA showed higher number of reads from HSPA1A compared to GFP (Figure 3B). We used Yodel peak-calling algorithm (Palmer et al., 2017) to analyse HSPA1A FLASH-seq data and identified 1087 binding sites of HSPA1A across the human transcriptome (Figure S2A and S2B). 300 of these RNA-binding sites mapped to 28S, 5.8S and 18S ribosomal RNA (rRNA), likely reflecting the interaction of HSPA1A with the ribosome-associated chaperone complex (Jaiswal et al., 2011). However, upon close inspection we found that the negative control GFP had similar RNA binding profile as HSPA1A on 28S, 5.8S and 18S rRNA suggesting a non-specific binding of these rRNA to tagged proteins or to beads during purification (Figure S2C). Of the remaining 787 binding sites specifically bound by HSPA1A, around 200 mapped to simple repeats of RNA, mainly poly(A) or poly(T) (Figure S2B). This may represent HSPA1A binding to poly(A) tail of mRNA or certain repetitive RNA classes that have poly(A) and poly(T) stretches. However, due to the low complexity of these sequences, it is difficult to attribute the binding of HSPA1A to mRNA or specific simple repeat-containing RNA.

**Figure 3.**
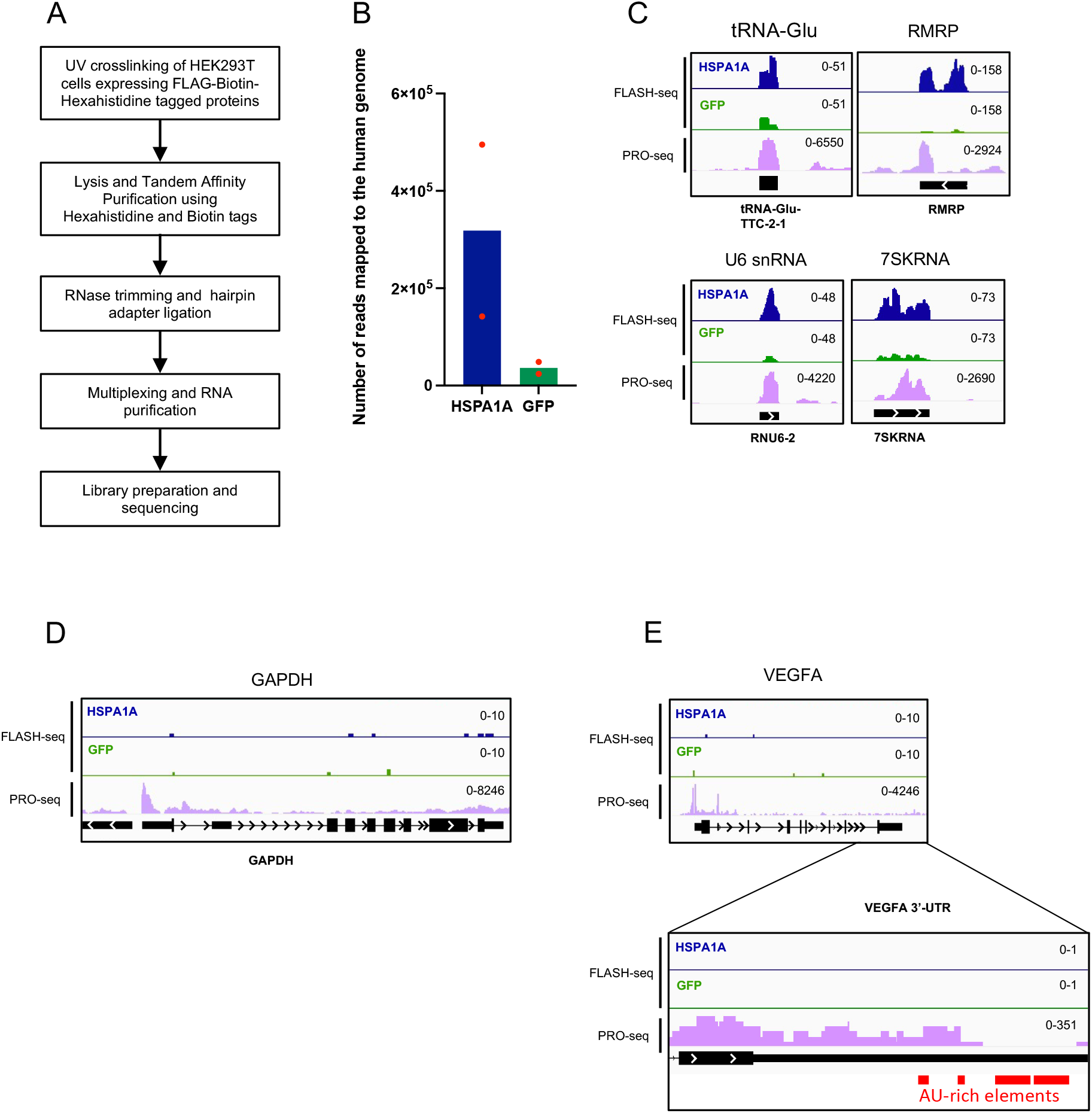
HSPA1A binds to non-coding RNAs. **(A)** Scheme of FLASH-seq workflow. **(B)** Number of reads mapped to the human genome obtained from FLASH-seq experiments performed using 3xFLAG-Biotin-Hexahistidine tagged HSPA1A or GFP expressing HEK293 T-Rex cell lines. The mean of two independent replicates is shown in the bar plot along with the values for each replicate as red dots. **(C-E)** Genome browser snapshots showing the distribution of HSPA1A- and GFP FLASH-seq reads over selected loci. Reads from Precision Run On sequencing (PRO-seq) dataset show nascent transcription from the corresponding genes. The inset in (E) represents the enlarged region corresponding to the 3’-UTR of the VEGFA gene where the position of AU-rich elements is highlighted by red boxes.

Majority of remaining RNA-binding sites of HSPA1A mapped to non-coding RNA (ncRNA) including small nuclear RNA (snRNA), tRNA, 5S rRNA and 7SK RNA (Figure S2A). A direct visualization of FLASH-seq reads on genome browser confirmed that HSPA1A binding was highly enriched as compared to GFP on these RNA species (Figure 3C, Figures S2D-G). However, the binding of HSPA1A was specific to a subset of ncRNA, since abundantly expressed long ncRNA such as XIST, NEAT1 and MALAT1 were not bound (Figure S2D). Furthermore, HSPA1A did not bind to mRNA such as GAPDH (Figure 3D), despite its high expression as measured by Precision Run-On sequencing (PRO-seq) in HEK293T cells (Aprile-Garcia et al., 2019). We also did not find any binding of HSPA1A to previously reported AU-rich elements in the 3’-UTR of VEGFA gene (Kishor et al., 2013) (Figure 3E) or other genes encoding oncoproteins and inflammatory mediators (Figure S2H) (Henics et al., 1999; Kishor et al., 2013; Wilson et al., 2001). Thus, we have mapped the complete repertoire of RNA directly bound by a chaperone for the first time, allowing us to address the functional relevance of the widespread chaperone-RNA interactions using HSPA1A as an exemplar.

### HSPA1A is not involved in the maturation of tRNA precursors

One of the major classes of RNA bound by HSPA1A in our FLASH-seq experiment was tRNA (Figure 3C, S2A, S2E). The binding of HSPA1A to the non-coding RNA RPPH1 in addition to tRNA (Figures 4A) suggested the involvement of HSPA1A in tRNA processing driven by RPPH1. The non-coding RPPH1 RNA is the catalytic component of the RNase P ribonucleoprotein complex. RNase P is responsible for the maturation of precursor tRNAs (pre-tRNAs) by the endonucleolytic cleavage of the 5’ leader sequence of pre-tRNAs (Figure 4B) (Mann et al., 2003). We observed that HSPA1A/B co-immunoprecipitated with RNase P components RPP29, POP1 and RPP21 (Figure 4C), along with the RNA component RPPH1 (Figure 4A). To assess if HSPA1A plays a role in the assembly, disassembly or function of RNase P ribonucleoprotein complex, we used small-molecule inhibitor of HSP70 (Williamson et al., 2009), which targets ATPase activity of HSPA1A along with all other HSP70 paralogues expressed in cells. Inhibition of HSP70 did not affect the interaction of RNase P component RPP29 with HSPA1A/B (Figure 4C) or with RPPH1 RNA (Figure 4D). Furthermore, HSP70 inhibition did not change the interaction between three of the components of the RNase P, RPP29, RPP21 and POP1 as observed in co-immunoprecipitation experiments (Figure 4E), ruling out the role of HSP70 (including HSPA1A) in assembly of RNase P complex. To assess if RNase P activity might be regulated by HSP70, we measured the efficiency of pre-tRNA processing *in vitro,* in the presence of HSP70 inhibitor or supplementing the reaction with recombinant purified HSPA1A (Figure 4F). RNase P complex was immunoprecipitated using the tagged RPP29 component. The endonucleolytic cleavage of pre-tRNA mediated by the affinity purified RNase P was comparable to the reaction catalysed by the positive control M1 RNA, the bacterial RNase P homologue (Figure 4G, lanes 1-3 and Figure S3A). The efficiency of pre-tRNA processing was not affected by the inhibition of HSP70 during in the *in vitro* RNase P assay (Figure 4G, lanes 3-4 and Figure S3B), or by supplementing the reaction with recombinant purified HSPA1A (Figure 4H, lanes 3-4). Furthermore, HSPA1A overexpression in HEK293T cells prior to purification of RNase P did not significantly change the activity of RNase P complex (Figure 4I, lanes 3-4 and Figure S3C). Thus, increasing the amount of HSPA1A *in vitro* or in cells did not alter RNase P activity. Altogether our data indicate that HSPA1A interacts with RNase P but does not regulate RNase P assembly or function.

**Figure 4.**
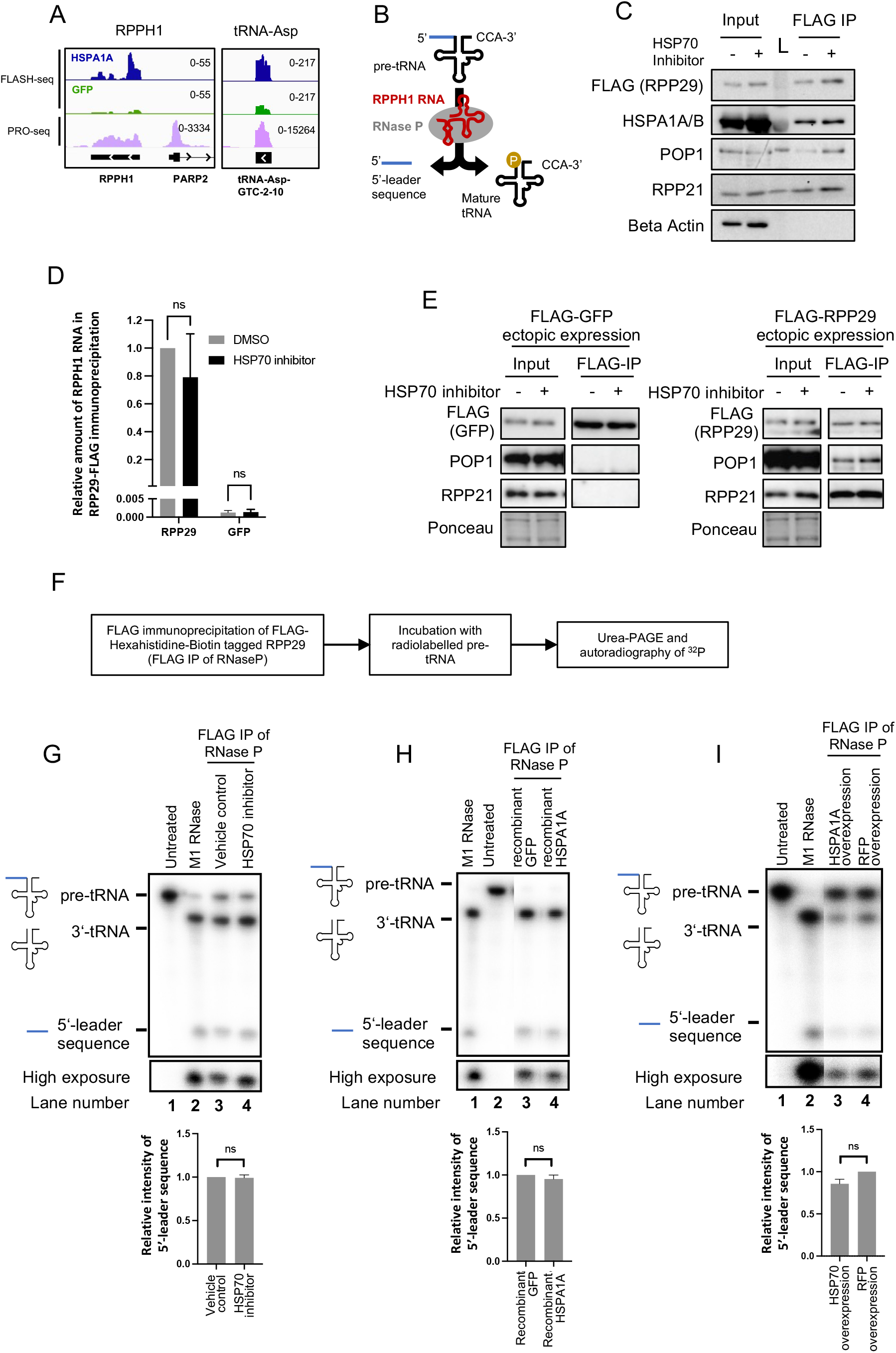
HSPA1A is not involved in the maturation of pre-tRNA. **(A)** Genome browser snapshots showing the distribution of HSPA1A and GFP FLASH-seq reads over genes encoding RPPH1 RNA and tRNA-Asp. Reads from Precision Run On sequencing (PRO-seq) show nascent transcription from the corresponding genes. **(B)** Schematic representation of RNase P-mediated endonucleolytic cleavage of the pre-tRNA 5’-leader sequences. **(C)** Western blot analysis of co-immunoprecipitation experiment using extracts of HEK293T cells expressing 3xFLAG-Biotin-Hexahistidine tagged RPP29 and treated with/without HSP70 inhibitor. Immunoprecipitation was performed with FLAG antibody. L denotes the protein molecular weight ladder. **(D)** Quantitative RT-PCR analysis of RPPH1 RNA pulled down in immunoprecipitation experiments using extracts from HEK293T cells treated with/without HSP70 inhibitor and expressing either 3xFLAG-Biotin-Hexahistidine tagged RPP29 or GFP. Immunoprecipitation was performed with FLAG antibody. The average of three independent replicates is shown; error bars represent S.E.M. Statistical analysis was performed using one sample t test (ns = non-significant). **(E)** Western blot analysis of co-immunoprecipitation experiment using extracts of HEK293T cells treated with/ without HSP70 inhibitor. HEK293T cells either expressed 3xFLAG-Biotin-Hexahistidine tagged GFP or RPP29. Immunoprecipitation was performed with FLAG antibody. **(F)** Experimental workflow of the pre-tRNA processing assay. **(G-I)** Autoradiograph of pre-tRNA processing assay (top) and quantification of the radioactive signal from the 5’-leader sequence (bar plots at the bottom). An average of five (G), three (H) and four (I) independent replicates is shown; error bars indicate S.E.M. Statistical analysis was performed using one sample t test (ns = non-significant). Bacterial M1 RNase was used as a positive control. RNase P was affinity purified from extracts of HEK293T cells expressing 3xFLAG-Biotin-Hexahistidine tagged RPP29. Pre-tRNA processing reactions were incubated in the presence of HSP70 inhibitor (G), recombinant HSPA1A (H) or by using affinity purified RNaseP from extracts of RFP- or HSPA1A-overexpressing HEK293T cells (I).

### HSPA1A binds chromatin at RNA Pol III transcribed loci

The identification of HSPA1A-bound RNA by FLASH-seq revealed an extensive binding of HSPA1A to RNA transcribed by RNA Polymerase III (Pol III). tRNA, RMRP, RPPH1, 7SK RNA are all products of Pol III transcription (Figures 3C, S2C, S2E-F). On the other hand, RNA Pol II transcripts such as mRNA and long ncRNA did not interact with HSPA1A (Figures 3D, 3E, S2D), with the exception of snRNAs (Figure S2G). 5S rRNA – the only ribosomal RNA that is transcribed by RNA Pol III – was bound by HSPA1A (Figure S2F). The other ribosomal RNAs that are transcribed by RNA Pol I showed a non-specific binding to HSPA1A in our FLASH-seq data (Figure S2C). RNA Pol III as a common source for most RNA interacting with HSPA1A suggested that HSPA1A may interact with RNA Pol III at chromatin and may bind RNA products co-transcriptionally. Indeed HSPA1A/B co-affinity purified with the tagged PolR3C (Figure 5A). Similarly, the endogenous RNA Pol III component PolR3A co-affinity purified with tagged HSPA1A (Figure 5B), confirming the interaction of HSP70 with the RNA Pol III machinery in cellular extracts. To further assess if HSPA1A could interact with RNA Pol III during transcription on genomic DNA, we mapped genome-wide binding sites of HSPA1A using chromatin immunoprecipitation followed by high-throughput sequencing (ChIP-seq). To compare the chromatin occupancy of HSPA1A with that of RNA Pol III, we identified RNA Pol III target genes which are active in HEK293T cells as inferred from the binding of RNA Pol III (Choquet et al., 2019). Strikingly, HSPA1A occupancy around transcription start sites (TSS) of active RNA Pol III-target genes was very similar to the occupancy of RNA Pol III (Figure 5C). RNA Pol III-target genes that are inactive in HEK293T cells and did not bind to RNA Pol III, did not show any signal of HSPA1A occupancy (Figure 5D). Visualizing HSPA1A ChIP-seq and FLASH-seq reads on genome browser confirmed the chromatin co-occupancy of HSPA1A and RNA Pol III at active genes as well as the binding of the corresponding RNA to HSPA1A at majority of loci (Figures 5E and S4A and S4B). The inactive genes were neither bound by HSPA1A nor by RNA Pol III at chromatin, and HSPA1A did not bind the corresponding non-coding RNA (Figure 5F).

**Figure 5.**
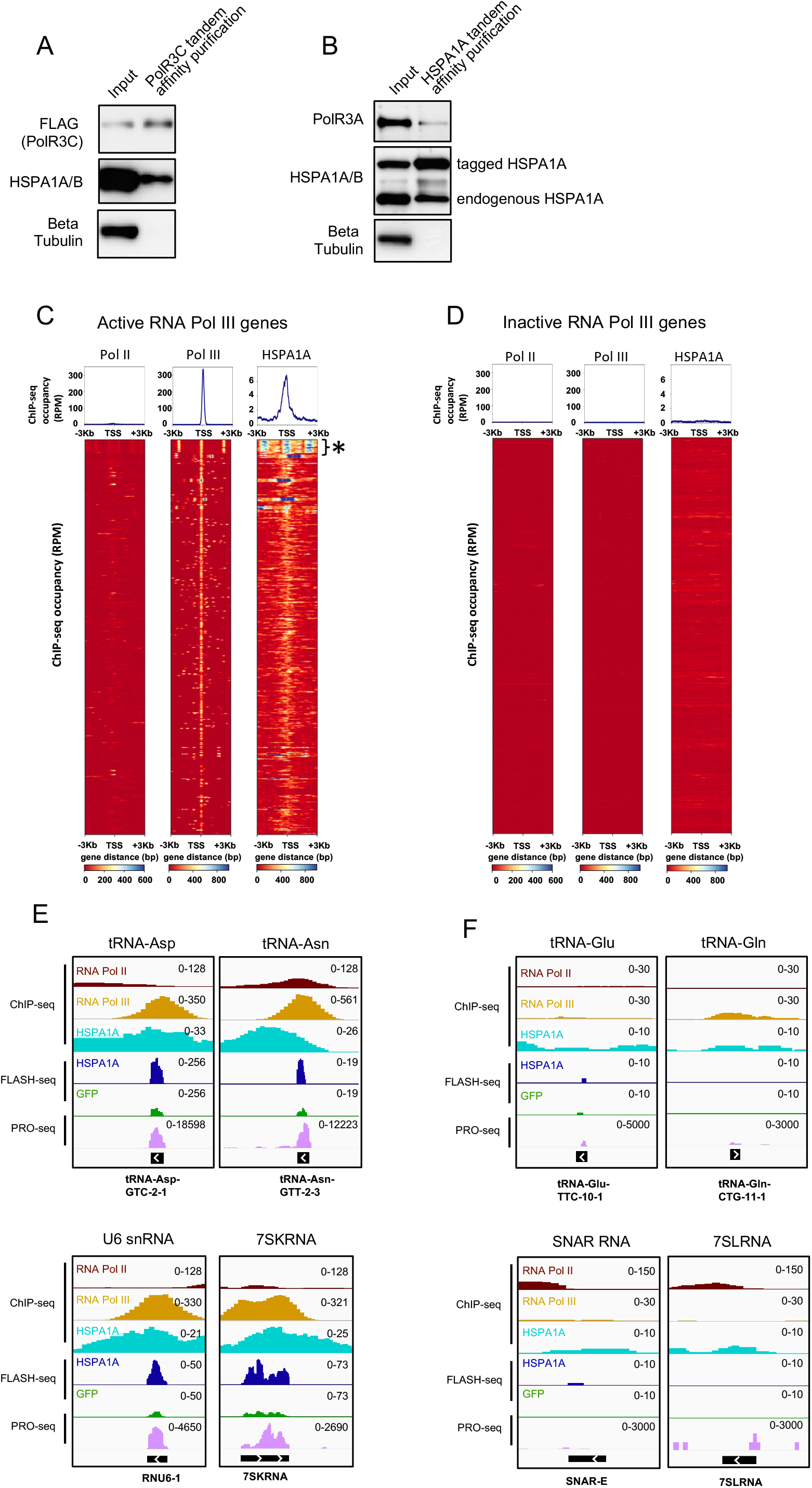
HSPA1A binds chromatin at RNA Pol III-transcribed loci. **(A, B)** Western blot analysis of co-immunoprecipitation experiments using extracts of HEK293T cells expressing either 3xFLAG-Biotin-Hexahistidine tagged POLR3C (A) or HSPA1A (B). Co-immunoprecipitation was done by tandem affinity purification using Hexahistidine and Biotin tags. **(C-D)** ChIP-seq analysis of RNA Pol II, RNA Pol III and 3xFLAG-Biotin-Hexahistidine tagged HSPA1A. Metaplots (top) and heatmaps (bottom) of indicated proteins binding around transcription start sites (TSSs) of 479 active (C) and 4105 inactive (D) RNA polymerase III transcribed genes. Y-axes indicate normalized ChIP-seq occupancy in reads per million (RPM). Color-scaled intensities are in units of RPM. * indicates the 5S-rRNA gene cluster composed of 2 Kb tandem repeats (see Figure S4A for a genome browser snapshot of 5S rRNA gene cluster). **(E-F)** Genome browser snapshots showing RNA Pol II, RNA Pol III and HSPA1A occupancy and the distribution of HSPA1A and GFP FLASH-seq reads over selected active (E) and inactive (F) genomic loci. Reads from Precision Run On sequencing (PRO-seq) show nascent transcription from the corresponding genes.

### HSPA1A regulates RNA Pol III transcription

The co-localization of HSPA1A and RNA Pol III at chromatin (Figure 5C-F, Figure S4) and direct interaction of HSPA1A with a majority of non-coding RNAs transcribed by RNA pol III (Figures 3C and Figure S2) suggested a functional link between HSPA1A and RNA pol III. To test the role of HSPA1A in RNA Pol III function, we performed Nuclear Run On (NRO) reactions (Figure 6A) as a quantitative measure of transcriptional activity of RNA Pol III in human cells (Roberts et al., 2015). We tested a subset of RNA Pol III genes which encoded the non-coding RNA bound by HSPA1A in our FLASH-seq data (Figures 3C, 4A and Figure S2). Inhibition of the ATPase activity of HSP70 during NRO reactions resulted in reduced transcription of RNA Pol III transcribed genes compared to vehicle treated control reactions (Figure 6B). NRO reactions use chromatin-assembled genomic DNA as template for transcription, reflecting a cellular role of HSP70 in RNA Pol III function. However, the effect of HSP70 inhibition could be indirect, for example, via the chaperone’s influence on chromatin structure or nucleosomes (Sawarkar and Paro, 2013) rather than the direct role of HSP70 in RNA Pol III activity. Also, HSP70 inhibitor targets all HSP70 isoforms, not just HSPA1A. We therefore employed an assay using a tRNA reporter plasmid as template for *in vitro* transcription (IVT) reactions instead of chromatin-assembled genomic DNA (Figure 6C). The reporter plasmid used in this assay contained the tRNA-Asp-GTC-2-10 gene, since both the gene locus and its transcript were bound by HSPA1A respectively in our ChIP-seq and FLASH-seq data (Figure S4B). Transcription of the tRNA-Asp reporter plasmid was reduced when HSP70 was inhibited in the reaction (Figure 6D). Importantly, supplementing the reaction with recombinant HSPA1A increased the transcriptional output (Figure 6E), confirming the role of HSPA1A in RNA Pol III function.

**Figure 6.**
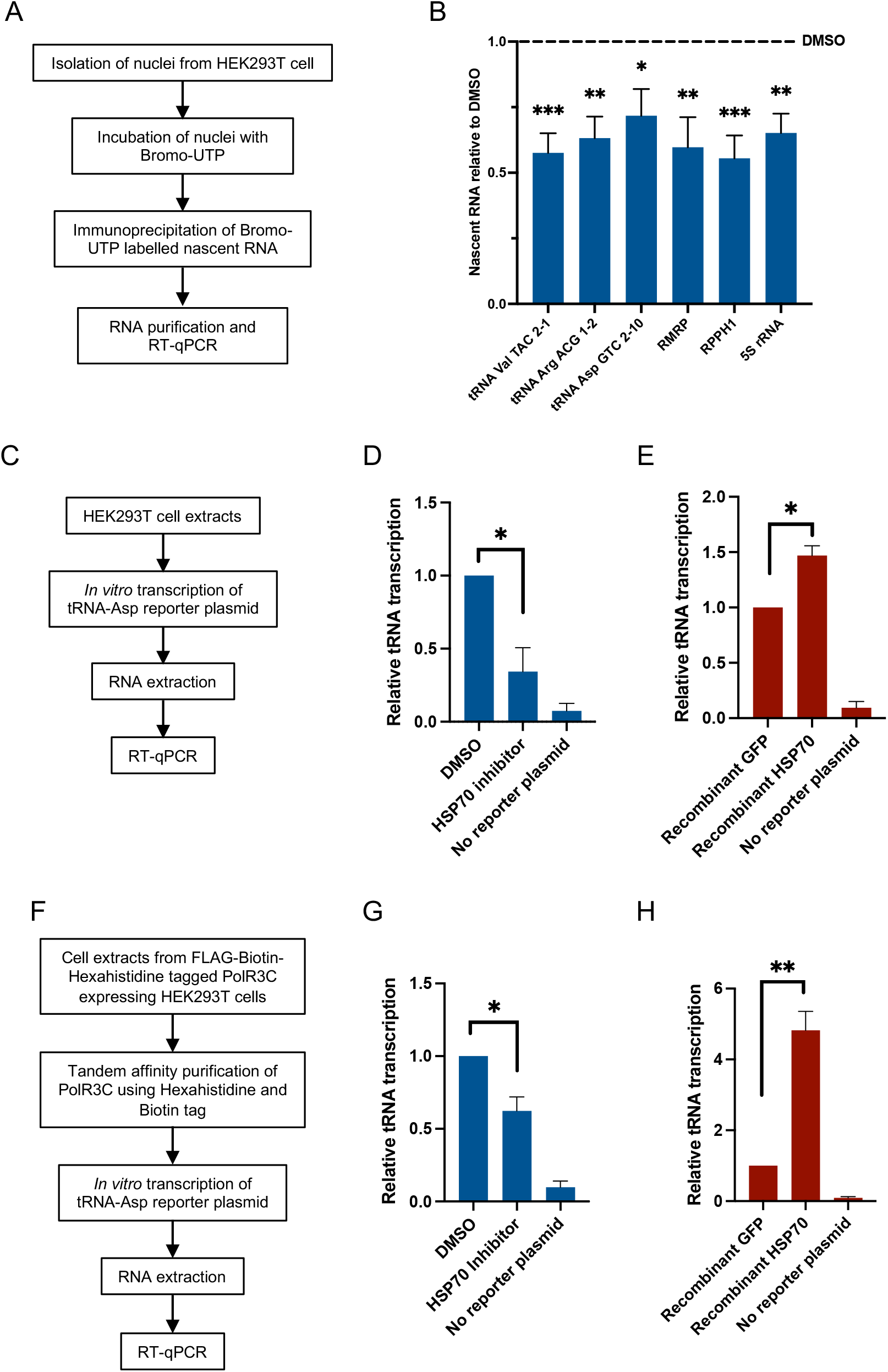
HSP70 regulates RNA polymerase III transcription. **(A)** Scheme of experimental workflow of nuclear run-on experiments. **(B)** Quantitative RT-PCR of indicated genes after nuclear run-on experiments performed in presence of DMSO or HSP70 inhibitor. Data are normalized over vehicle control and the mean of ten replicates is shown. Error bars indicate S.E.M. Statistical analysis was performed using one sample t test. Asterisks denote P values (*\**, 0.05; *\*\**, 0.01 and *\*\*\**, 0.001). **(C)** Scheme of experimental workflow of *in vitro* transcription assays performed using HEK293T cellular extracts and tRNA-Asp-GTC-2-10 reporter plasmid. **(D-E)** Quantitative RT-PCR analyses of reporter tRNA transcription reaction detailed in (C). HEK293T cellular extracts treated with HSP70 inhibitor or vehicle control (D) or supplemented with recombinant HSPA1A or recombinant GFP (E) were used in the reaction. Data are normalized over vehicle control or GFP control respectively and the average of three and four (E) independent replicates is shown. Error bars indicate S.E.M.. Statistical analysis was performed using one sample t test. Asterisks denote P value (*\**, 0.05). **(F)** Experimental workflow of *in vitro* transcription assays performed using tRNA-Asp-GTC-2-10 reporter plasmid and tandem-affinity-purified PolR3C from HEK293T expressing 3xFLAG-Biotin-Hexahistidine tagged PolR3C. **(G-H)** Quantitative RT-PCR analyses of reporter tRNA transcription reaction detailed in (F). Transcription reaction performed using tandem affinity purified 3xFLAG-Biotin-Hexahistidine tagged POLR3C. Reactions were performed in presence of HSP70 inhibitor or vehicle control (G), or recombinant HSPA1A or recombinant GFP (H). Data are normalized over vehicle or GFP control respectively and the mean of five (G) and four (H) independent replicates is shown. Error bars indicate S.E.M. Statistical analysis was performed using one sample t test. Asterisks denote P values (*\**, 0.05; *\*\**, 0.01).

Total cell extracts used in the *in vitro* transcription reactions contain several proteins and RNA which could indirectly contribute to the effect of HSP70 inhibition on RNA Pol III transcription. Hence, we employed a refined system wherein tandem affinity purified RNA Pol III was used instead of total cell extracts for *in vitro* transcription of tRNA reporter plasmid (Figure 6F). Transcription driven by purified RNA Pol III was reduced by the presence of HSP70 inhibitor in the reaction (Figure 6G), indicating that the endogenous HSP70 copurified with RNA Pol III (Figures 5A and 5B) has an active role in modulating the transcriptional output even in a purified system. Furthermore, supplementing the reaction with recombinant HSPA1A increased the transcription of the reporter plasmid more than 4-fold compared to reaction supplemented with recombinant GFP (Figure 6H), confirming that HSPA1A acts as a positive regulator of RNA Pol III transcription.

In summary, our work provides the evidence for the interaction of a large number of chaperones with RNA. Using HSPA1A as an example, this study shows that the chaperone binds to specific regions of chromatin co-occupied by RNA Pol III and interacts with non-coding RNAs transcribed by RNA Pol III. Finally, refined biochemical assays suggest that HSPA1A facilitates the transcriptional activity of RNA Pol III.

## DISCUSSION

The unexpected widespread interaction between chaperones and RNA reported here raises two fundamental questions. First, how do chaperones contribute to RNA function, biogenesis, or stability? As highlighted in this pioneering study, the unbiased identification of chaperone-bound RNA in intact cells will likely provide clues to the breadth of chaperone function in specific aspects of RNA biology. Changes in chaperone-RNA interactions during ageing and in pathological conditions will be an important area of study with ramifications on therapeutic targeting of chaperones. In this context, the role of RNA, protein disordered domains, and molecular chaperones in biomolecular condensates formation both in healthy and diseased cells may be an exciting avenue to investigate (Ganassi et al., 2016; Kato et al., 2012; Yang et al., 2020). The second fundamental question raised by chaperone-RNA interaction is how RNA contributes to the function of chaperones in maintaining the cellular proteome? Biochemical assays of chaperone activity carried out with addition of different RNA species will be key to understanding the molecular contribution of protein-RNA interactions to chaperone function. Furthermore, structural insights into chaperone-RNA interaction will be useful in identifying chaperone mutants that lack RNA-binding capacity for functional studies. The role of RNA in regulating the chaperone function during proteotoxic stress will be an important aspect to study. Chaperones such as CCT and HSP90 are known to play a role in ribonucleoprotein assembly by directly binding protein members of the complexes (Iwasaki et al., 2010; Zhao et al., 2008). It is likely that chaperones directly interact with RNA constituents of ribonucleoprotein complexes during the assembly process. In this regard, it is worth noting that we observed a substantial binding of HSP70 to small nuclear RNAs (snRNAs; Figure 3C, S2A and S2G), which are part of the spliceosome complex. HSP70 is shown to be a direct interactor of spliceosome components in both human and fly cells (Herold et al., 2009) and is involved in the recovery of mRNA splicing after heat inactivation (Vogel et al., 1995). Thus, the interaction of HSP70 with snRNA might be explained by a direct interaction between HSP70 and the mRNA splicing machinery in the nucleus.

The identification of ER-resident chaperones binding to RNA is surprising as the ER lumen is devoid of RNA. The ER-resident chaperones may localize briefly outside the ER-lumen such as during trafficking of PDIA3 and Calnexin to the cell surface in cancer cells (Ros et al., 2020), providing an opportunity to crosslink with RNA associated with the cytosolic surface of the ER membrane.

Our inability to recapitulate HSP70 binding to intergenic rRNA and 3’ UTRs of mRNA may stem from the fact that we identified HSP70-RNA interactions that take place in intact cells, unlike previous studies that used cellular extracts or purified recombinant proteins to document HSP70-RNA interactions (Audas et al., 2012; Henics et al., 1999; Kishor et al., 2013). Nonetheless, the identification of poly(A) sequences in our dataset (Figure S2A and S2B) suggests that HSP70 may directly bind to poly(A) tails of mRNA in agreement with previous studies on HSP70-mRNA interactions. While the yeast HSP70 homologue Ssb1 has been described to directly interact with rRNA (Gumiero et al., 2016), the interaction of the human HSPA1A with ribosomal RNA appears to be non-specific as our negative control GFP also bound exactly the same sequences on rRNA (Figure S2C). Nonetheless, HSP70 interaction with RNA reported in this study by protein-RNA crosslinking in intact cells is further corroborated by the orthogonal demonstration of chromatin binding of HSP70 precisely at the site of expression of HSP70-bound RNA (Figure 5 and Figure S4). HSP70 may regulate RNA pol III activity by chaperoning the different phases of the transcription cycle, namely initiation, elongation, termination and reinitiation (Turowski and Tollervey, 2016). Thus, detailed structural and reconstitution experiments will be required to dissect the molecular mechanism by which HSP70-RNA interaction regulates RNA pol III.

In summary, our study provides the first global view of chaperone-RNA interaction in human cells paving an avenue to investigate how RNA and proteostasis may be functionally coupled in healthy and diseased cells.

## ACKNOWLEDGEMENTS

We would like to thank A. Akhtar, T. Aktas, V. Dezi, M. Elzek, V. Hilgers, I. A. Ilik, N. Jarrous, M. Pizzinga, E. Trompouki for reagents, scientific discussions and technical assistance. We also thank proteomics- and deep sequencing facilities of Max Planck Institute of Immunobiology and Epigenetics, Freiburg, Germany. This work was supported by funds from the Max Planck Society (Germany), the Medical Research Council (UK) and the European Research Council (ERC) Consolidator grant.

## AUTHOR CONTRIBUTIONS

S.L. and R.S. conceived the project and designed the experiments. S.L. and A.S. conducted the experiments. L.T., F.A.G., A.S, and P.R. helped with the experiments. B.H. and A.H.R. carried out computational analyses. S.L. and R.S. wrote the manuscript with inputs from A.E.W, R.S. supervised the study.

## DECLARATION OF INTERESTS

The authors declare no competing interests.

## DATA AND CODE AVAILABILITY

All data supporting the findings of the current study are available within the article and its Supplemental Information files or from the corresponding authors upon reasonable request. All deep-sequencing data generated in this study are deposited in GEO and are available under accession number GSE191245. The following secure token has been created to allow review of record GSE191245 while it remains in private status: **klgrkcmildullsx**

## FIGURE LEGENDS

**Figure S1 (related to Figure 1).**
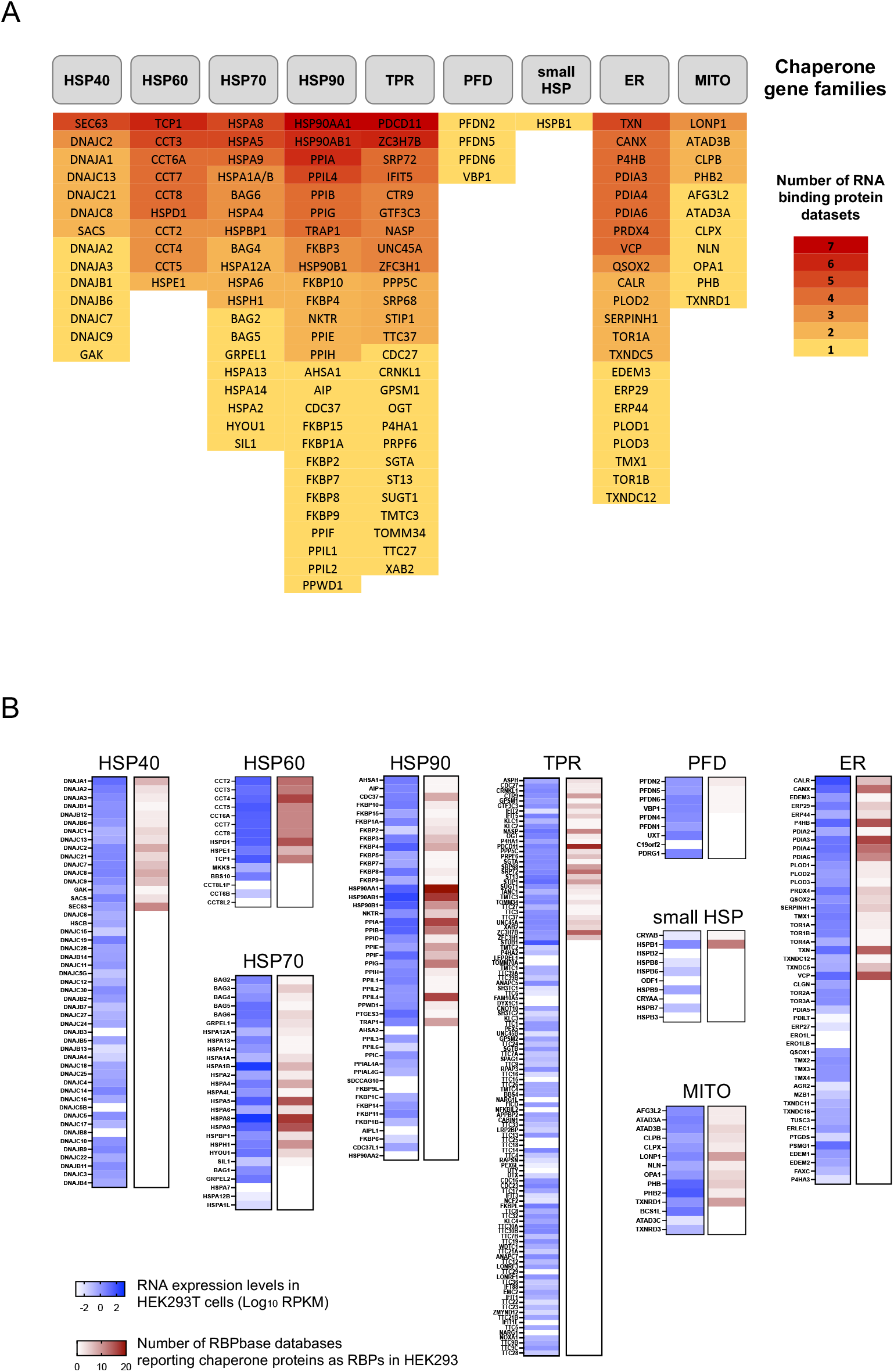
**(A)** List of RNA-binding chaperome proteins identified in independent datasets using HEK293 cells. The proteins are sorted into chaperone families (TPR: tetratricopeptide repeat (TPR)-domain-containing protein; PFD: prefoldin; ER: endoplasmic reticulum specific proteins; MITO: mitochondria specific proteins). The boxes containing the protein name are color-coded and sorted in descending order based on the number of independent datasets in which they were identified as RNA-binding proteins. Number of datasets reporting HSPA1A and HSPA1B are merged since the two proteins have 100% amino acid identity and are not distinguishable by mass spectrometry analyses. **(B)** Comparison of chaperone genes expression (RNA-seq data, blue heatmaps) and the number of datasets reporting chaperome proteins as RNA-binding proteins (red heatmaps) in HEK293 cells. Proteins are grouped according to the chaperone family.

**Figure S2 (related to Figure 3).**
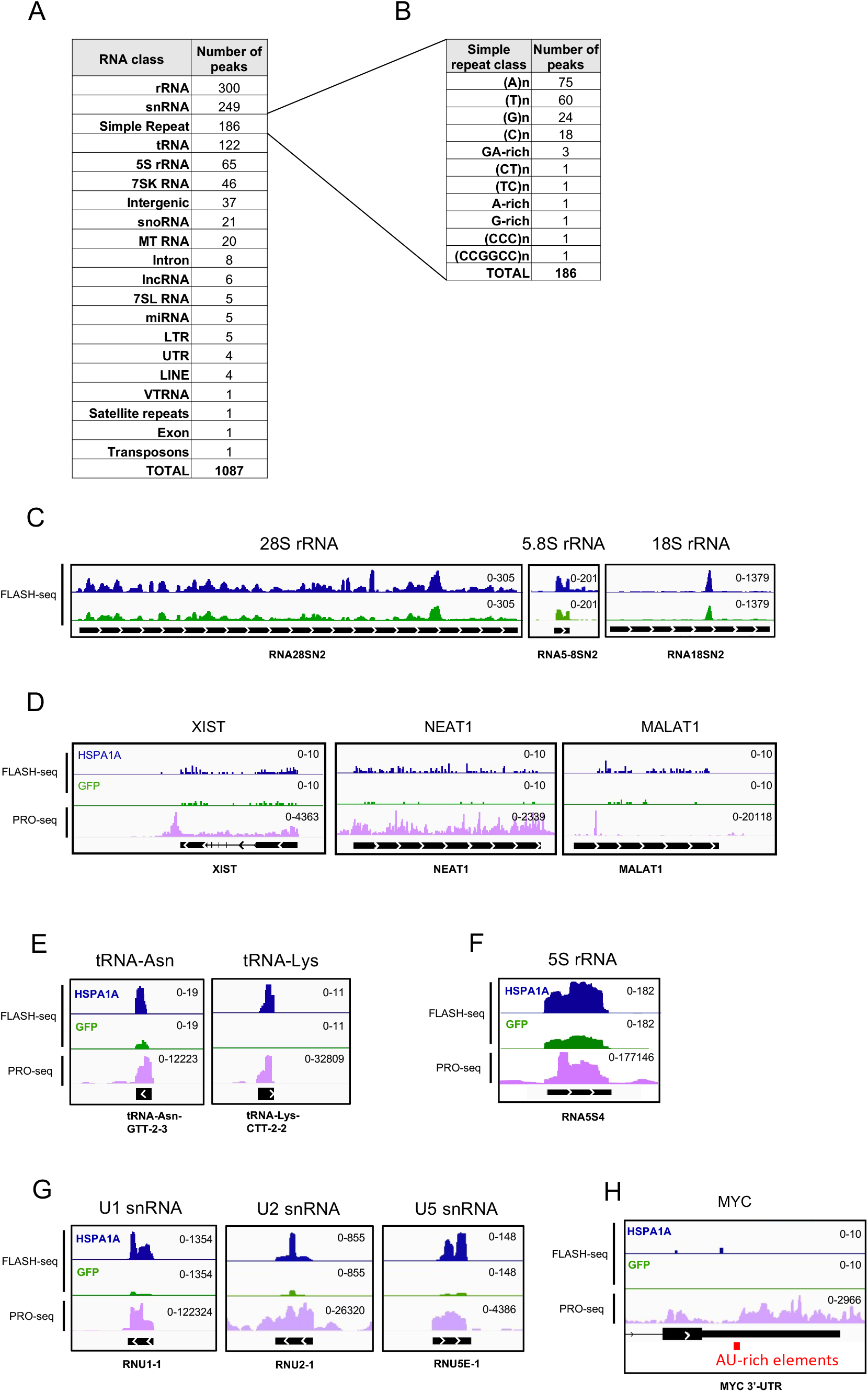
**(A)** The distribution of peaks obtained in HSPA1A FLASH-seq across different RNA classes. **(B)** Details of Simple Repeat RNA class peaks obtained in HSPA1A FLASH-seq experiment shown in (A). **(C)** Genome browser snapshots showing the distribution of HSPA1A and GFP FLASH-seq reads over 28S, 5.8S and 18S ribosomal RNA genes. **(D-H)** Genome browser snapshots showing the distribution of HSPA1A and GFP FLASH-seq reads over selected loci. Reads from Precision Run On sequencing (PRO-seq) show nascent transcription from the corresponding genes. The region corresponding to the 3’-UTR of the MYC gene is shown in (H) and the position of AU-rich elements is highlighted by red boxes.

**Figure S3 (related to Figure 4).**
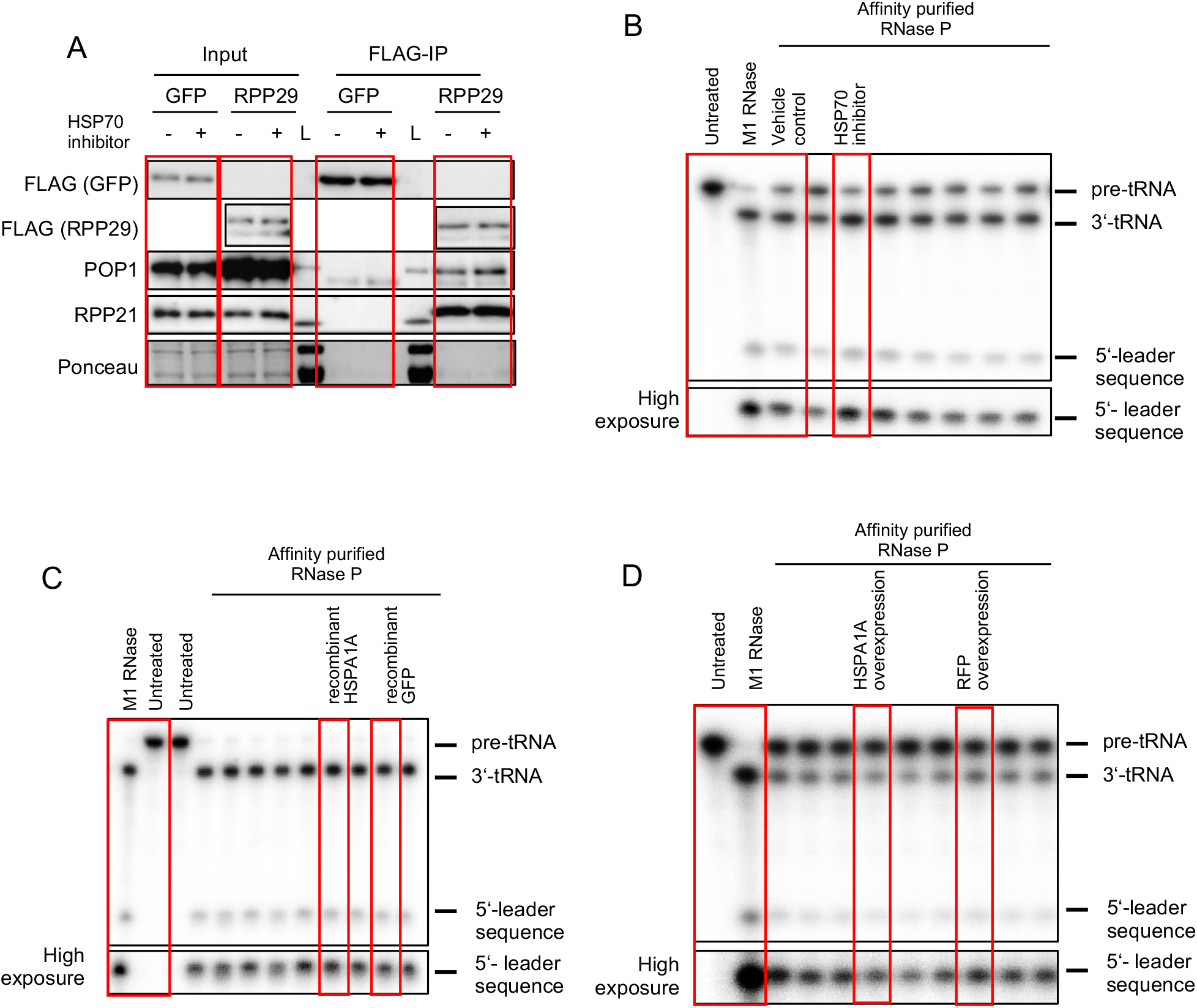
**(A)** Full size Western blot of co-immunoprecipitation experiment shown in Figure 4E. **(B)** Full size autoradiograph of pre-tRNA processing assay shown in Figure 4G. Red boxes indicate data shown in Figure 4G. **(C)** Full size autoradiograph of pre-tRNA processing assay shown in Figure 4H. Red boxes indicate data shown in Figure 4H. **(D)** Full size autoradiograph of pre-tRNA processing assay shown in Figure 4I. Red boxes indicate data shown in Figure 4I.

**Figure S4 (related to Figure 5).**
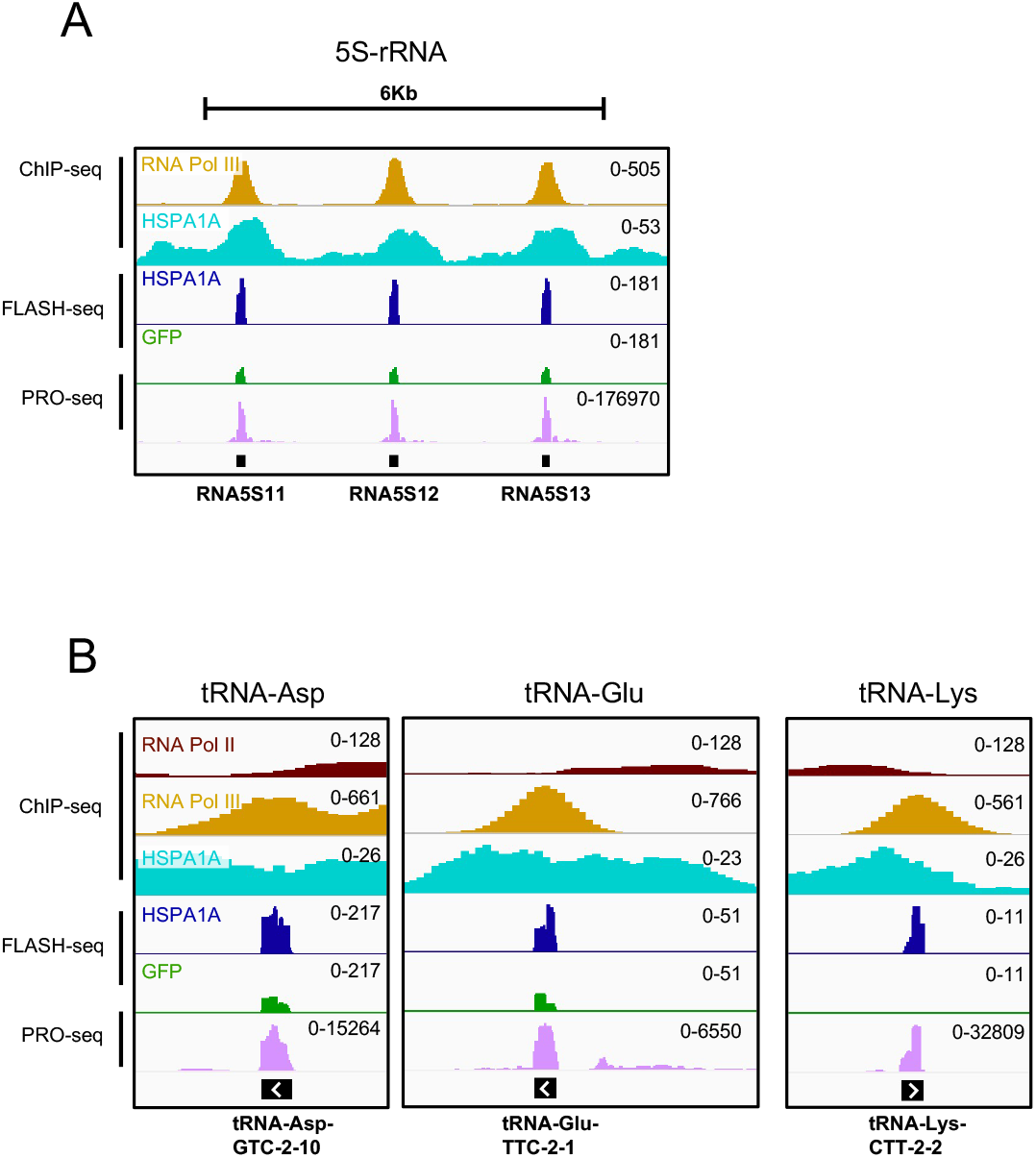
**(A)** Genome browser snapshot showing RNA Pol II, RNA Pol III and HSPA1A occupancy and the distribution of HSPA1A and GFP FLASH-seq reads over a representative 6kb region that includes three 5S-rRNA loci and reflects the occupancy of RNA Pol III and HSPA1A observed for the genes in the * cluster of the heatmaps shown in Figure 5C. **(B)** Genome browser snapshots showing RNA Pol II, RNA Pol III and HSPA1A occupancy and the distribution of HSPA1A and GFP FLASH-seq reads over tRNA genes.

## MATERIALS AND METHODS

### Cell culture

HEK293 Flp-In T-Rex cells were cultured in DMEM high glucose medium supplemented with 10% FBS (Sigma-Aldrich F0804), 2mM L-glutamine (Sigma-Aldrich, G7513) and 1% penicillin-streptomycin (Sigma-Aldrich, P4333). Cells were kept at 37 °C in 5% CO_2_ incubator. All cell lines were routinely checked for mycoplasma contamination by PCR.

### Plasmid cloning

For generation of clones for human proteins, hORFs in pDONOR vector were purchased from BIOSS, University of Freiburg and cloned into pCDNA5-FRT/To vector with a C-term 3xFLAG-HBH tag (kind gift from Ibrahim Avsar Ilik, Max Plank Institute, Freiburg) using Gateway LR clonase II enzyme kit (Life Technologies 11791020). pSupS1 and pJA2 M1 RNA were kindly provided by Prof. Nayef Jarrous (The Hebrew University of Jerusalem, Jerusalem)

### Plasmid transfection

HEK293T Flp-In T-Rex cells were transfected using calcium phosphate protocol. To generate stable expression cell lines Flp-In T-Rex HEK293 were grown in 100 μg/ml Zeocin (Invitrogen, R25001) for at least a week before transfection. Stable positive clones were selected for at least 2 weeks in 15 μg/ml blasticidin S (Carl Roth CP14.1) and 100 μg/ml hygromycin B (ThermoFisher Scientific, 10687010). Expression of chaperone protein was induced with 1 μg/ml tetracycline (Sigma-Aldrich, T7660).

### Tandem Affinity Purification

Cell were washed in 1xPBS and collected by scraping in NLB buffer (1xPBS; 300mM NaCl; 1% Triton-X100; 0.1% Tween-20). Cell lysis was obtained by pulsed ultrasonication (5 cycles, 30” On, 30” Off, at High intensity using Diagenode Bioruptor 300). Lysates were incubated 10 minutes at 4°C with Dynabeads™ His-Tag Isolation (Invitrogen, 10103D). After one wash with NLB buffer, proteins were eluted with 250mM imidazole in NLB buffer and incubated for 45 minutes rotating at 4°C with Dynabeads™ MyOne™ Streptavidin C1 (Invitrogen, 65001). After one wash in HSB buffer (50mM Tris-HCl, pH 7.4; 1M NaCl; 1% Igepal; 0.1% SDS; 0.5% Na-deoxycholate; 1mM EDTA) and one wash in NDB buffer (50mM Tris-HCl, pH 7.4; 100mM NaCl; 0.1% Tween-20). Beads containing purified proteins were resuspended in NDB buffer and used for PNK assay and FLASH-seq.

### Polynucleotide kinase (PNK) assay

Cells were grown as monolayers and irradiated on ice with 200mJ/cm^2^ of 254nm UV light. After washing in 1xPBS, cells were collected by scraping in NLB buffer (1xPBS; 300mM NaCl; 1% Triton-X100; 0.1% Tween-20) and lysed by pulsed ultrasonication (5 cycles, 30” On, 30” Off, at High intensity using Diagenode Bioruptor 300). Cell lysates were subjected to tandem affinity purification as described above. To produce PNK accessible 5’-ends, “minus RNase” samples were incubated with 0.05U/µl of RNase I at 37°C for one minute. For a complete digestion of RNA “plus RNase” samples were treated with NDB buffer containing 2U/µl of RNase I (Invitrogen, AM2294) and 0.02µg/µl of RNase A (Thermo Scientific, EN0531) at 37 °C for 15 minutes. After RNase treatment, beads were washed once with HSB, once with NDB buffer and twice with PNK buffer (20mM Tris-HCl pH 7.4; 100mM MgCl2; 0.2% Tween-20). 1/10 of the affinity purified complexes were eluted for 10 minutes at 70°C in 1x LDS buffer (Invitrogen, NP0007) containing 25mM DTT and analysed by SDS PAGE and Western blot. The rest of the affinity purified complexes were radiolabelled with 0.1 µCi/µl γ-^32^P ATP by T4 polynucleotide kinase (1U/µl) in PNK buffer for 30 minutes at 37 °C shaking at 1100rpm. Beads were washed twice with PNK buffer and protein-RNA complexes were eluted for 10 minutes at 70°C in 1x LDS buffer (Invitrogen, NP0007) containing 25mM DTT. Samples were analysed by SDS PAGE and autoradiography.

### Chromatin immunoprecipitation and sequencing

Cells were fixed using 2mM Di(N-succinimidyl) glutarate (DSG; Sigma-Aldrich, 80424-50MG-F) for 40 minutes in 1xPBS at room temperature and 1% methanol-free formaldehyde (Thermo Scientific, 28906) was added in the last 10 minutes, followed by 5 min blocking in 0.125 M glycine. Cells were washed twice with ice-cold PBS. The cell pellet was resuspended in Farnham buffer (5 mM PIPES, pH 8; 85 mM KCl; 0.5% Igepal). Cell suspensions were sonicated for 3 minutes in 1 ml Covaris tubes (Covaris, 520130) using Covaris S220 with the following settings: peak power = 75; duty factor = 2; cycles/ burst = 200. Isolated nuclei were washed with Farnham buffer and suspended in shearing buffer (10 mM Tris-HCl, pH 8; 0.1% SDS; 1 mM EDTA). Chromatin was sheared by sonication in 1-ml Covaris tubes using the following settings: peak power = 140; duty factor = 5; cycles/burst = 200, time = 25–30 min. Debris were removed by centrifugation at 16000 x g. A DNA fragment–size distribution of 200–600 bp was considered as ideal chromatin for ChIP. Chromatin was diluted 1:1 with IP buffer (10 mM Tris-HCl, pH 8; 100 mM NaCl, 1 mM EDTA, 0.5 mM EGTA, 0.1% sodium deoxycholate, 0.1% N-lauroylsarcosine) to achieve a final 0.05% SDS concentration. Good quality chromatin (200 μg) was used for immunoprecipitation. Protein G magnetic beads (Life Technologies, 10004D) were incubated (rotated) with 5–10 μ g of FLAG antibody (Sigma-Aldrich, F1804-200UG) for 6 hours at 4 °C. This bead–antibody complex was then incubated overnight at 4 °C with sheared chromatin. An aliquot of chromatin was saved as input DNA. Beads were washed and DNA–protein complexes were eluted from the beads by heating at 65 °C in elution buffer (50 mM Tris-HCl, pH 8.0, 10 mM EDTA and 1% SDS). Crosslinking was reversed for 6 h at 70 °C and samples were treated with 200 μg/ml RNase A (Thermo Scientific, EN0531) and 200 μg/ml proteinaseK (Sigma-Aldrich, P2308). ChIP DNA was purified with phenol–chloroform extraction and ethanol precipitation. Some 2–5 ng of immunoprecipitated DNA was used for library preparation for next-generation sequencing. Sequencing libraries were prepared using the NEBNext Ultra II DNA Library Prep kit for Illumina (NEB E7645S). Library size distribution was monitored by capillary electrophoresis (Agilent 2100 Bioanalyzer, High Sensitivity DNA Chips (Agilent, 5067–4626)). Libraries were sequenced paired end on NextSeq500 or NovaSeq6000 instruments (Illumina).

### FLASH sequencing

FLASH-seq protocol was applied as described in Aktaş et al., 2017 and Maticzka et al., 2018. Briefly, doxycycline-induced HEK293T Flp-In T-Rex cells were rinsed in 1xPBS and crosslinked with 0.2mJ/cm2 UV-C irradiation. Crosslinked cells were collected by scraping and pelleted by centrifugation. Cells were resuspended in NLB buffer and lysed and by water bath-sonication. Target protein of interest were affinity purified in tandem with nickel-charged paramagnetic beads and streptavidin-coupled paramagnetic beads against the tagged protein of interest as described above (Tandem affinity purification). To trim the crosslinked RNA, the beads were resuspended with 1 mL of NDB (50 mM Tris-Cl, pH 7.4; 100 mM NaCl, 0.1% Tween-20), to which 2 µL of TURBO DNase (AM2238, Thermo Fisher Scientific) and 10 µL of diluted RNaseI (1:2000–1:8000 dilution in NDB from 100 U/µL stock (AM2294, Thermo Fisher Scientific)) were added. The solution was incubated at 37 °C for 3 min and immediately transferred on ice. Dephosphorylation of cyclic phosphate groups was carried out with T4 PNK (10 U/µL, M0201, NEB) in a low pH buffer (25 mM MES (2-(N-morpholino)ethanesulfonic acid), pH 6.0; 50 mM NaCl; 10mM MgCl2; 0.1% Tween-20; 20U RNasin (Thermo fisher, AM2694); 10 U PNK; 20 min at 37 °C). Custom-made, barcoded 3’-adapters were ligated using T4 RNA Ligase 1, for 1 hr at 25°C. Custom FLASH adapters contained two barcodes and random nucleotides adjacent to the 3’-adapters according to the pattern NNBBNTTTTTTNN (N: random tag nucleotide, T: tag nucleotide, B: RY-space tag nucleotide). Random tags were used to merge PCR-duplicates, regular tags were used to specify the pulldown condition, and semi-random RY-space tags were used to distinguish the biological replicates (RR: replicate A, YY: replicate B, R: purine, Y: pyrimidine). Excess adapters were washed away, and RNA was isolated with Proteinase K treatment and column purification (Zymo DNA Clean and Concentrator). Isolated RNA was reverse transcribed with SuperScript III (Invitrogen, 18080093) and treated with RNaseH (M0297S NEB). cDNA was column-purified and circularized with CircLigase (Epicentre/Illumina, CL9021K) for 16hrs. Circularized cDNA was directly PCR amplified, quantified with Qubit/Bioanalyzer and sequenced on Illumina NextSeq 500 or NextSeq 3000 in paired-end mode.

### Bioinformatic analysis

FLASH-seq data were demultiplexed using Flexbar, version 3.3 (Roehr et al., 2017). For each sample, UMIs were extracted using UMITools, version 0.5.1 (Smith et al., 2017) followed by adapter removal with TrimGalore, version 0.4.4 [http://www.bioinformatics.babraham.ac.uk/projects/trim_galore/]. Potential readthroughs into the barcode and UMI region were removed by clipping the last 13 bases from the 3’ ends of first mate reads. Paired-end reads were merged using bbmerge from bbmap, version 37.54 [https://jgi.doe.gov/data-and-tools/bbtools/bb-tools-user-guide/]. The demultiplexed and processed reads were mapped to the human genome build hg38 using STAR, version 2.6.0b (Dobin et al., 2013) with the addition of the parameters “--outFilterMultimapNmax 150 -- outFilterScoreMinOverLread 0 --outFilterMatchNminOverLread 0 --outFilterMatchNmin 0 -- alignEndsType EndToEnd --alignIntronMax 100000”. UMITools, version 0.5.1 (Smith et al., 2017) was used to combine duplicated reads into individual crosslinking events. For each FLASH sample, enriched regions were identified using YODEL (Palmer et al., 2017). Resulting peak regions were annotated using homer annotatePeaks.pl (Heinz et al., 2010). Bigwig files were generated using deeptools2 (Ramírez et al., 2016). Yodel peaks were cleared from alternative chromosomes.

### Whole cell extract preparation

Cells were harvested by scraping in ice-cold 1xPBS and washed once in Buffer A (10 mM KCl, 1.5 mM MgCl2, 0.5 mM DTT, 10 mM Hepes, pH 7.9). Cells were resuspended in one packed cell volume of Buffer A, incubated on ice for 10 minutes and passed through a 19G needle 10 times. After adjusting the salt concentration to 200mM KCl the cells were passed through a 21G needle. Cell lysis was obtained by pulsed ultrasonication (10 cycles, 30” On, 30” Off, at High intensity using Diagenode Bioruptor 300), complete lysis was monitored by phase contrast microscopy. Cellular debris were pelleted at 20000 x g for 20 minutes at 4°C. The supernatant was used for immunoprecipitation or for *in vitro* transcription experiments.

### Immunoprecipitation

Whole cell extracts were prepared as described above. Immunoprecipitation was carried out using FLAG-coupled agarose beads (Millipore, A2220) in IP buffer (20mM Tris-HCl pH 7.5; 100mM KCl; 0.5mM DTT; 0.2mM EDTA; 20% glycerol). Protein content was measured by BCA assay (Thermo Scientific 23225). 250-500µg of protein extracts were incubated rotating at 4°C for 16 hours with 20-40 µl of IP buffer pre-washed FLAG-coupled agarose beads. After incubation, beads were washed three times with TNET-150 buffer (20mM Tris-HCl pH7.5; 150 mM NaCl; 0.1 mM EDTA; 1mM beta-mercaptoethanol; 0.01% v/v Triton-X100) and three times with PA buffer (50mM Tris-HCl pH7.5; 0.1M NH_4_Cl; 10mM MgCl_2_). Immunoprecipitated proteins were eluted in 20-40µl of 1x TNET and 1x PA buffer containing 300ng/µl of FLAG peptide for 10 minutes at room temperature. Eluted proteins were used in the RNase P assay or Western blot.

### RNase P assay

RNase P assay was performed as described in Mann et al., 2003. Briefly, 5µl of the 3xFLAG- HBH-RPP29 co-immunoprecipitated RNaseP holoenzyme or the M1 RNA control were incubated in 20µl 1xTNET 1xPA buffer supplemented with 1mM DTT, 1X Protease inhibitor (Sigma-Aldrich, 1187358000) and 10U of RNase inhibitor (Ambion, AM2694) at 37°C for 1 hour. Reactions were supplemented either with 20µM VER155008 HSP70 inhibitor (Sigma-Aldrich, SML0271-5MG), 50nM recombinant HSPA1A (Cayman Chemicals, 22739) or 50nM recombinant GFP (Sino Biological, 13105-S07E). The yeast pre-tRNASer (pSupS1), was transcribed *in vitro* by T7 RNA polymerase in the presence of [alpha-^32^P] CTP, purified on a 7M urea 5% polyacrylamide gel, and was used at a final concentration of 500 nM (ca. 2,000 cpm per reaction). Gels were exposed to phosphorimager screen at -20°C for 16 hours.

### In vitro transcription

In vitro transcription assays were performed using whole cell extracts or RNA Pol III holoenzyme co-purified with 3xFLAG-HBH-PolR3C tandem affinity purification. IVT reaction were carried as described in Baer et al., 1990. Briefly, 25 µl of whole cell extracts or 20 µl of RNA Pol III-containing streptavidin paramagnetic beads were incubated in a final volume of 50µl for 30-60 minutes in 1X transcription buffer (12 mM Tris-HCl pH7.9; 5 mM MgCl_2_; 80 mM KCl, 0.5 mM DTT; 20 mM creatine phosphate; 0.5 mM each of ATP, UTP, CTP, GTP; 20U of RNase inhibitor) containing 500 ng of pUC-tRNA-Tyr reporter plasmid. Reactions were supplemented either with 20 µM Pifithrin µ HSP70 inhibitor (Sigma-Aldrich P0122-5MG), 140 µM DMSO (Sigma-Aldrich, 276855), 50 nM recombinant HSPA1A (Cayman Chemicals, 22739) or 50 nM recombinant GFP (Sino Biological, 13105-S07E). IVT reaction were terminated by adding 10 volumes of Trizol reagent (Invitrogen, 15596026).

### Nuclear Run-on

Nuclear Run-on assays were performed as described in Roberts et al., 2015. Briefly, nuclei were obtained from cells incubated in NP-40 lysis buffer (10 mM Tris-HCl, pH 7.4, 10 mM NaCl, 3 mM MgCl2 and 0.5% (vol/vol) NP-40). Run-on was carried out incubating nuclei in transcription buffer (10 mM Tris-HCl, pH 8.3, 2.5 mM MgCl2, 150 mM KCl, 2 mM DTT, 1mM each of ATP, CTP, GTP; 0.5 mM of each UTP and Bromo-UTP; 20U of RNase inhibitor). Run-on reactions were carried out in the presence of 20µM VER155008 HSP70 inhibitor (Sigma-Aldrich, SML0271-5MG) or 140 µM DMSO (Sigma-Aldrich, 276855) and incubated at 30 °C for 40 minutes. Reactions were stopped with 10 volumes of Trizol reagent (Invitrogen, 15596026) containing 0.01pg/µl of bromouridine-labelled *in vitro* transcribed renilla luciferase used as positive spike-in control, or 0.01 pg/µl firefly luciferase (Promega, L4561) used as negative spike-in control.

### RNA purification and reverse transcription

RNA purification was performed using Trizol reagent (Invitrogen, 15596026) according to the manufacturer instructions. RT reaction was done using Takara PrimeScript RT reaction kit (RR047A) and qPCRs were done using Takara TB Green Premix Ex-Taq (RR420L) according to the manufacturer instructions.

### RNA sequencing

RNA purification was performed using Trizol reagent (Invitrogen, 15596026) according to the manufacturer instructions. TruSeq stranded Total RNA (Gold), Illumina was used for the construction of sequencing libraries.

## Notes

### Competing Interest Statement

The authors have declared no competing interest.

